# The effects of action-based predictions in early visual cortex

**DOI:** 10.1101/2025.08.28.672792

**Authors:** Bianca M. van Kemenade, Lars F. Muckli

## Abstract

During voluntary movement, predictions about the sensory consequences of an action typically result in reduced sensory sensitivity. The forward model theory proposes that this reduced sensitivity is due to neural suppression or cancellation of sensory action outcomes. However, recently this theory has been challenged by three alternative theories: the pre-activation account, sharpening, and the opposing processes theory. In this fMRI study, we directly tested and compared these four theories using univariate and multivariate analyses both prior to and during stimulus presentation. Participants performed a visual orientation discrimination task on two sequentially presented gratings, which were either presented automatically (passive condition) or triggered by their own button press (active condition). Auditory cues at trial onset indicated the overall grating orientation, followed by a preparatory phase in which participants anticipated the upcoming stimuli. During this phase, the predicted stimulus orientation was decodable from early visual cortex activity in both active and passive conditions at levels significantly above chance, indicating pre-activation of the predicted orientation, but with no difference between active and passive conditions. BOLD responses did not emerge earlier in active conditions, arguing against the pre-activation theory. During stimulus presentation, actively generated stimuli elicited larger BOLD responses than passively presented ones, contradicting the forward model theory, which predicts overall response suppression. Decoding accuracy did not differ between conditions, inconsistent with the sharpening hypothesis, which predicts enhanced neural precision for actively generated stimuli. Instead, our findings align most closely with the opposing processes theory, which posits pre-activation in both conditions. However, the stronger BOLD response for actively generated stimuli is not predicted by any existing theory, suggesting that additional mechanisms—such as heightened attention or motor-related enhancement—may contribute to sensory processing during self-initiated actions.

## Introduction

Our actions produce sensory outcomes, contributing substantially to the constant stream of sensory input that we encounter every day. It was proposed decades ago that we suppress sensory consequences of our own action using predictions generated by an internal forward model (Wolpert et al., 1995). According to this theory, this internal forward model uses the efference copy, a copy of the motor command, to generate predictions about the sensory outcomes of our action (Sperry, 1950; von Holst & Mittelstaedt, 1950). These predictions are then used to cancel out sensory action consequences, or generate prediction errors in the case of a mismatch between prediction and sensory feedback. This mechanism is thought to aid us in guiding our actions, learn new motor skills, save resources, and distinguish self from other (S. Blakemore & Frith, 2003; Wolpert & Flanagan, 2001). In line with this idea are numerous studies that show reduced neural activity elicited by self-generated stimuli compared to identical but externally generated stimuli (S.-J. Blakemore et al., 1998; Lubinus et al., 2022; Martikainen et al., 2005; Shergill et al., 2013; Straube et al., 2017). For many years, this theory has shaped our thinking about sensorimotor mechanisms. However, in recent years this theory has been challenged by three alternative theories. First of all, the preactivation account posits that suppression of self-generated stimuli is not due to a true dampening of neural activity, but rather a pre-activation of sensory areas (Roussel et al., 2013). The idea is that due to this pre-activation, the baseline activity prior to stimulus onset is increased, making it harder to distinguish the actual stimulus activity from the baseline activity, leading to an apparent suppression during stimulus presentation. However, evidence for this account using neuroimaging techniques is lacking. Secondly, studies on sensory predictions have suggested that the apparent suppression induced by sensory expectations, so called “expectation suppression” (Summerfield et al., 2008), may instead be due to a sharpening of the underlying neuronal populations. This neural sharpening may resemble suppression when using univariate analysis, but can be detected using multivariate pattern analysis showing better decoding of expected stimuli (Kok et al., 2012). It has been proposed that action also leads to neural sharpening (Yon et al., 2018), but to our knowledge no-one has directly compared active with passive conditions using this approach to confirm that the effects of action on neural activity can be specifically attributed to neural sharpening. Lastly, the recently proposed „opposing process theory” suggests that action-based predictions and general predictions about our environment are based on the same mechanism and involve different time scales (Press et al., 2019). According to this theory, any prediction about an upcoming event firstly pre-activates neurons tuned to the expected features. During stimulus presentation, “cancellation” may occur in the form of boosting the unexpected event compared to the expected event, but only when the unexpected event is likely to be informative. This theory unites the fields of action-based and general sensory predictions, but as it was proposed only very recently, the support for this theory is still preliminary. Evidence for pre-activation of neurons tuned to expected features exists in the perception literature (Kok et al., 2014, 2017), but evidence for neural pre-activation for action-based predictions, as well as a supposed similarity between these two types of predictions, is lacking.

In this fMRI study, we combined univariate and multivariate analysis techniques both prior to and during stimulus presentation in order to directly test and compare all four theories. This strategy allowed us to test for changes in both overall neural amplitude and representational content during both prediction and perception phases (see Table 1). If action cancels out sensory outcomes through dampening as the forward model theory suggests, we would expect both reduced neural activity and reduced decoding accuracy for self-generated stimuli. If action instead sharpens neural responses, we would expect reduced neural activity but increased decoding accuracy for self-generated stimuli. If action modulates neural activity through pre-activation, we would expect increased activity and increased decoding accuracy during preparation, and suppressed neural activity and decreased decoding accuracy during stimulus presentation. Lastly, if action-based and general sensory predictions follow the same principles as outlined in the opposing process theory, we would expect a pre-activation of expected units through significant above-chance decoding of the upcoming stimulus during preparation, with no differences between self-generated and externally generated conditions. During stimulus presentation, we would expect no differences in activity or decoding accuracy, at least when using highly predictable stimuli.

**Table 1.** Four models’ hypotheses about activity level and decodability of visual features during the preparation phase and in response to stimulus presentation. Marked in blue are unique predictions.

| Tab. 1 | Preparatory activity | Preparatory decoding | Stimulus activity | Stimulus decoding |
| --- | --- | --- | --- | --- |
| Forward Model |  |  | A- < P | A- < P |
| Sharpening |  |  | A- < P | A+ > P |
| Pre-activation | A+ > P | A+ > P | A- < P | A- < P |
| Opposing process | A = P | A+ = P+ | A = P | A+ = P+ |

**Table 2.**
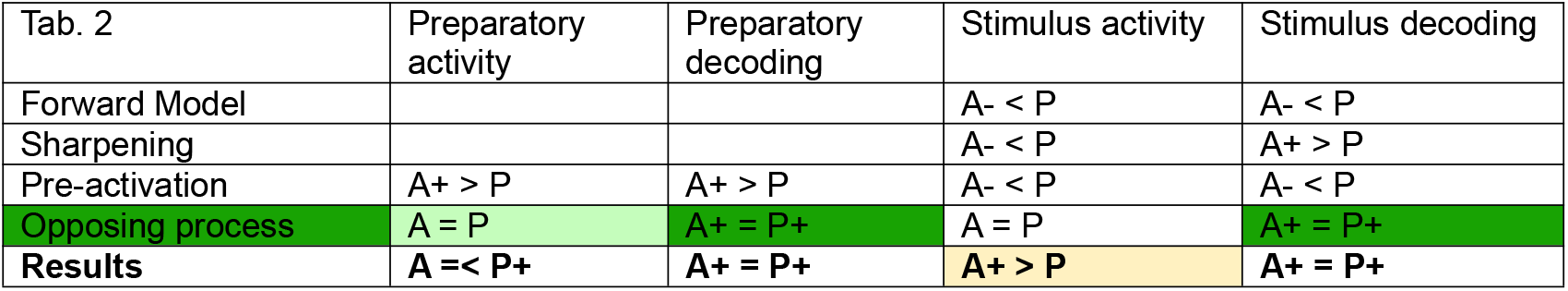
Results compared with the hypotheses from each theory. The above chance decoding during the preparatory and stimulus phases, with no differences between active and passive conditions, match the predictions made by the opposing process theory. The univariate results during the preparatory phase partly match the opposing process predictions. Only the enhancement during the stimulus phase does not match any prediction.

## Results

Participants performed both a behavioural and an fMRI experiment, that used the same stimuli and experimental paradigm (see Fig. 1). Participants had to perform an orientation judgment task on pairs of gratings that they either caused themselves to appear (active blocks) or that appeared automatically (passive blocks). Each trial started with an auditory cue that predicted with 100% validity the overall orientation of two consecutively presented gratings (45° leftward or 45° rightward). There was a preparation phase of eight seconds between the auditory cue and the active/passive stimulus generation, during which only a fixation dot was presented on the screen, but participants already had a prediction about the upcoming stimuli due to the cue. We performed our fMRI analyses for this preparation phase and for the stimulus phase separately. In the behavioural experiment, the preparation phase after the cue was reduced to three seconds.

**Fig. 1.**
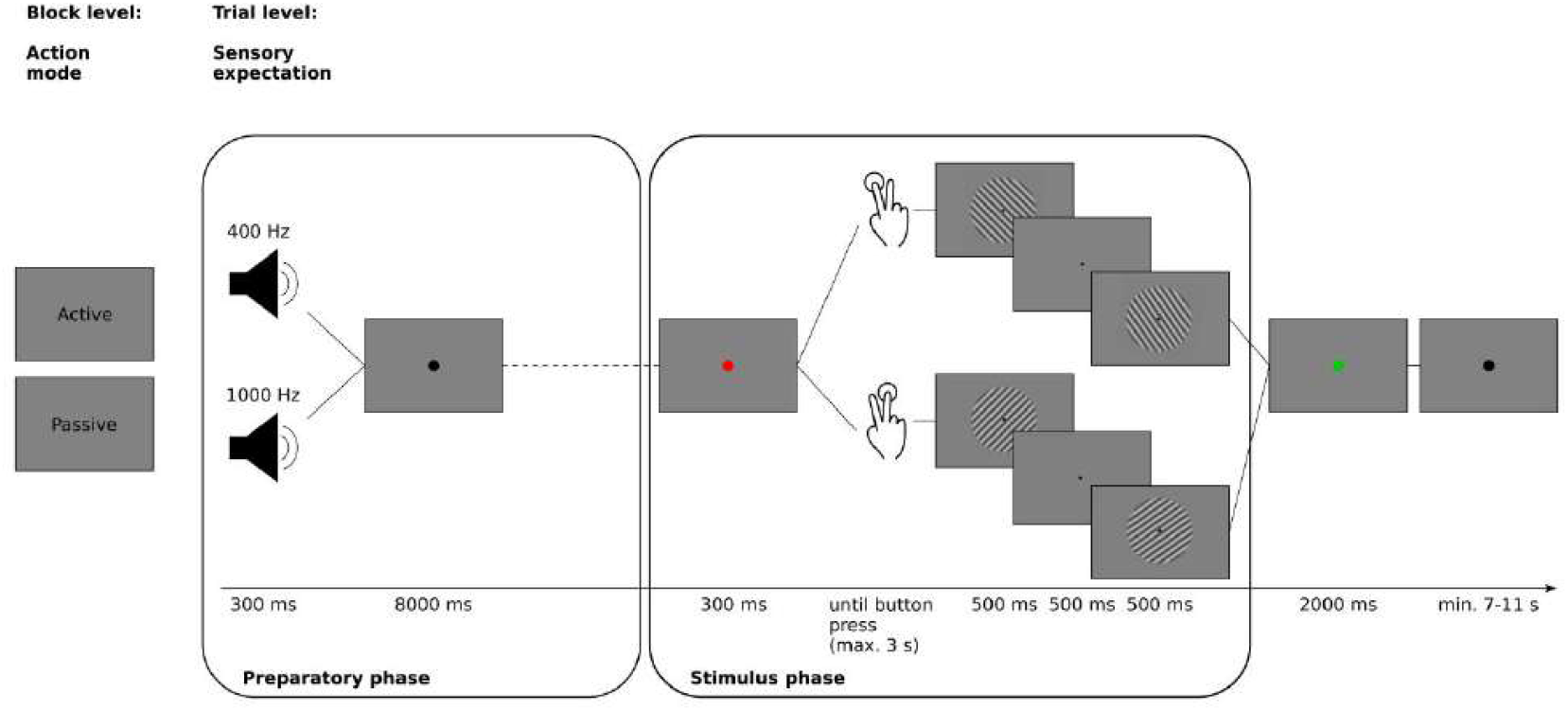
Experimental paradigm. Active and passive conditions were presented in separate blocks. Each trial started with a predictive cue indicating the overall orientation of the upcoming gratings and informing participants which button to press in the active condition. After the cue, a preparatory phase started during which only a fixation dot was shown. Then the fixation dot briefly turned red, which meant that participants had to now press a button with their index or middle finger to elicit the presentation of the gratings (active condition) or had to wait for the gratings to be presented automatically (passive condition). The two gratings differed slightly in their orientation (+/-0.5-6.5°), and participants were asked to judge whether the second grating was tilted clockwise or counterclockwise compared to the first one. They had to report their answer when the dot turned green after grating offset. Any time that remained from the button press window in the active condition to elicit grating presentation was added to the ITI (7-11 s, mean 9 s). This means that trials had on average the same duration (24,1 s). The mapping between cue frequency, button, and grating orientation was counterbalanced across participants.

### Behavioural effects

We fitted psychometric functions to the data from the behavioural experiment for active and passive conditions separately and derived thresholds and slopes. Furthermore, we calculated the overall accuracy. Paired-samples t-tests testing for differences between active and passive conditions revealed no significant differences, neither for thresholds (t(27)=0.209, p=0.836), nor slopes (t(27)=-0.216, p=0.831) or overall accuracy (t(27)=0.310, p=0.759), as displayed in Fig. 2.

**Fig. 2.**
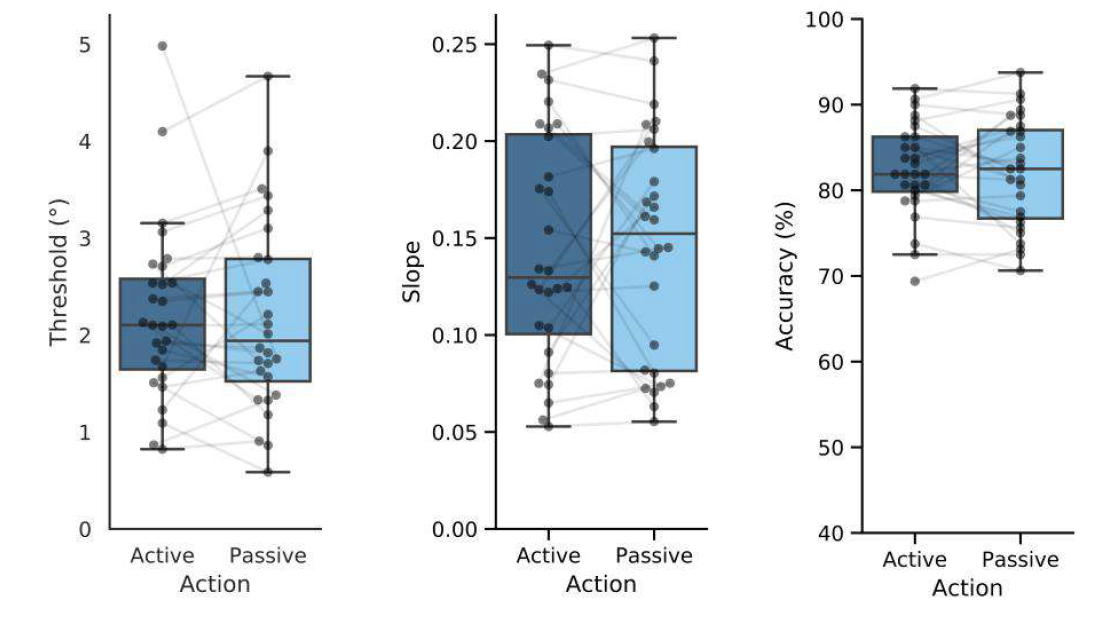
Behavioural results. Boxplots showing the thresholds and slopes derived from psychometric functions for each condition, as well as the overall accuracy.

We applied the same approach to the behavioural data obtained during the fMRI experiment. Replicating the findings from the behavioural experiment, we found no significant differences between active and passive conditions, neither for thresholds (t(25)=0.613, p=0.546), nor slopes (t(25)=0.155, p=0.878) or overall accuracy (t(25)=0.059, p=0.954), see Fig. 3.

**Fig. 3.**
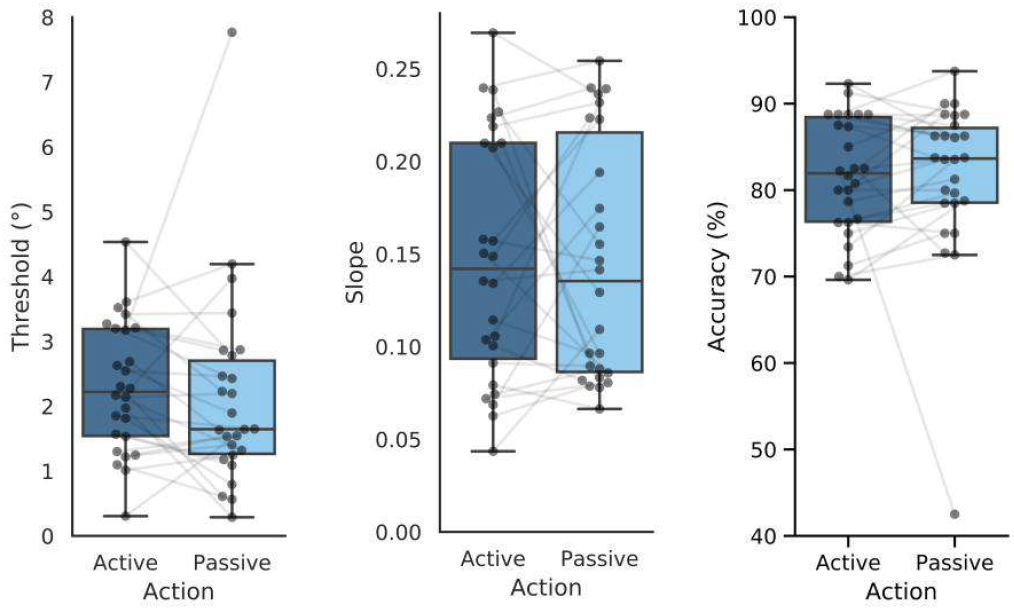
Behavioural results from the fMRI experiment. Boxplots showing the thresholds and slopes derived from psychometric functions for each condition, as well as the overall accuracy.

### Univariate analyses

Self-generated stimuli evoked more activity than externally generated stimuli in early visual cortex (t(25) = −4.463, p = 0.00015). During preparation, no significant differences between active and passive conditions were found (t(25) = −1.392, p = 0.176), see Fig. 4. The results were comparable across all ROIs (see Supplementary Material).

**Fig. 4.**
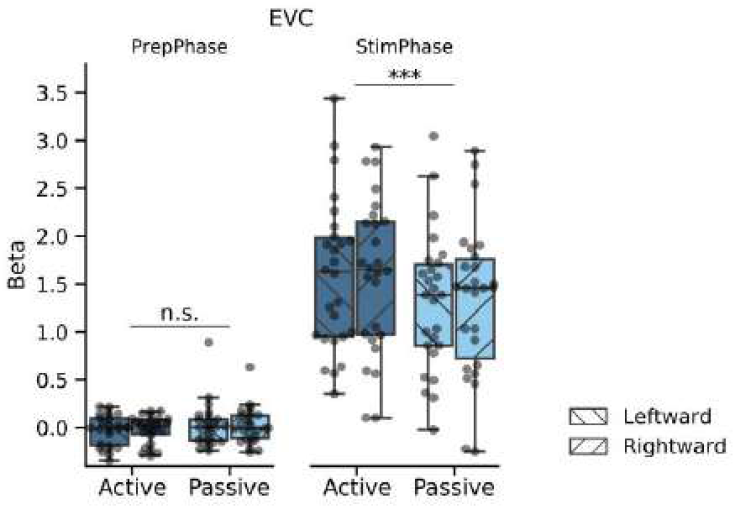
Univariate results. We performed paired t-tests to test for differences between active and passive conditions. Boxplots show the extracted beta values from early visual cortex (EVC). n.s. = not significant, *** = p < 0.001.

Next, in order to test whether there were any differences in the timing of the BOLD response, we performed a deconvolution GLM. As can be seen in Fig. 5, responses did not differ in their timing. Instead, amplitude differences were found in the late preparatory phase, with more sustained activity in the passive condition (V3: p = 0.0018; all other p < 0.001), and in the stimulus phase, with a larger response in the active condition (all p > 0.001). Responses did not differ in amplitude in the early preparatory phase (EVC: t(25) = −1.109, p = 0.267; V1: t(25) = −0.724, p = 0.469; V2: t(25) = −1.699, p = 0.089; V3: t(25) = −1.870, p = 0.062).

**Fig. 5.**
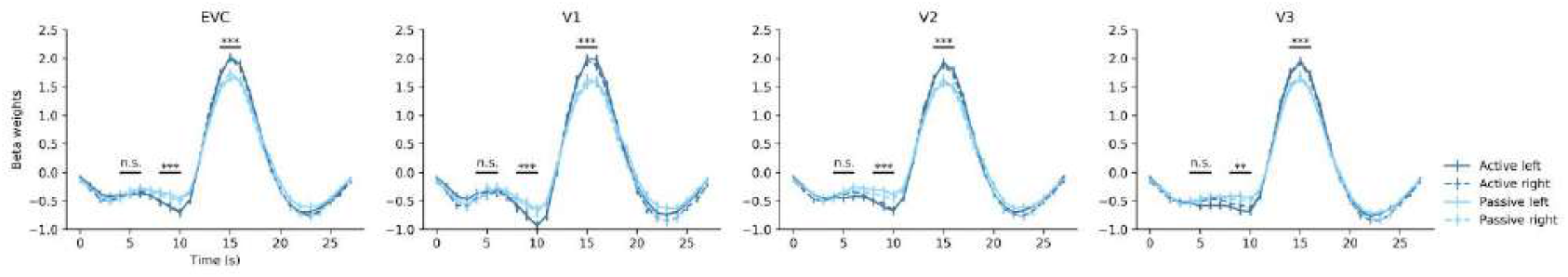
Results from deconvolution GLM. Time courses are plotted for each condition, time-locked to the start of a trial (auditory cue). Error bars represent the standard deviation. n.s. = not significant, ** = p < 0.01, *** = p < 0.001.

Our analysis of eye tracking data showed that the differences between active and passive conditions cannot be explained by differences in fixation behaviour, neither during the preparatory phase (t(13)=-1.345, p = 0.202) nor during the stimulus phase (t(13)=-1.752, p = 0.103).

### Multivariate analyses

Fig. 6 illustrates the results of our multivariate analyses. The orientation of the gratings could be decoded significantly above chance during the stimulus phase in all ROIs (active conditions: EVC 71.2%, V1 66.7%, V2 70.7%, V3 65.9%; passive conditions: EVC 74.5%, V1 70.8%, V2 70.8%, V3 67.8%; all p < 0.001), as was to be expected based on previous studies (Kamitani & Tong, 2005). Importantly, the orientation of upcoming, not yet presented stimuli in the preparatory phase could also be decoded significantly above chance, again in all visual areas (active conditions: EVC 54.5%, V1 54.4%, V2 54.1%, V3 52.0%; passive conditions: EVC 56.2%, V1 56.3%, V2 56.3%, V3 54.8%, V3 active conditions p = 0.025; all other p < 0.001). There were no significant differences between active and passive conditions in any visual area, neither during the preparatory nor the stimulus phase (PrepPhase: all p > 0.195; StimPhase: EVC t(25) = −1.723, p = 0.097, V1 t(25) = −1.840, p = 0.078, V2 t(25) = −0.022, p = 0.983, V3 t(25) = −1.247, p = 0.224).

**Fig. 6.**
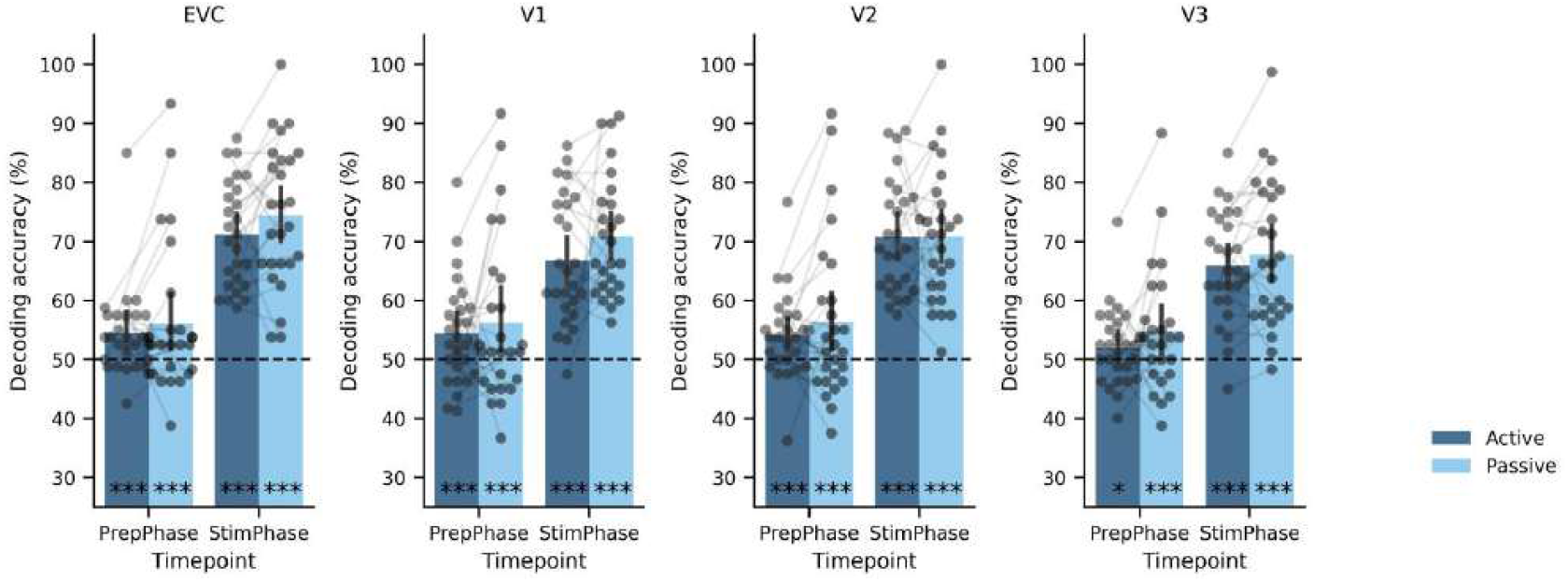
Results from multivariate analysis. Decoding accuracy is plotted for each single participant. Error bars represent the 95% confidence interval. Chance level is indicated by the dotted line at 50%. * = p < 0.05, *** = p < 0.001.

In order to test for similarities between active and passive conditions, we performed cross-classifications (Fig. 7). Training the classifier on active trials and testing on passive trials (and vice versa) revealed very similar results, showing significant above-chance decoding in both the preparatory and stimulus phase, indicating the classifier could generalise across conditions (Training on active and testing on passive trials, PrepPhase: 55.4%; StimPhase: 73.4%. Training on passive and testing on active trials, PrepPhase: 54.1%, StimPhase: 71.8%; all p < 0.001). In contrast, training on the preparatory phase and testing on the stimulus phase (and vice versa) showed chance-level decoding for both active and passive conditions, suggesting that the underlying patterns differed between these time points (Training on the PrepPhase and testing on the StimPhase, active conditions: 48.9%, p = 0.869, passive conditions: 48.8%, p = 0.892. Training on the StimPhase and testing on the PrepPhase, active conditions: 49.1%, p = 0.941, passive conditions: 49.8%, 0.620). Results were comparable across all ROIs (see Supplementary Material).

**Fig. 7.**
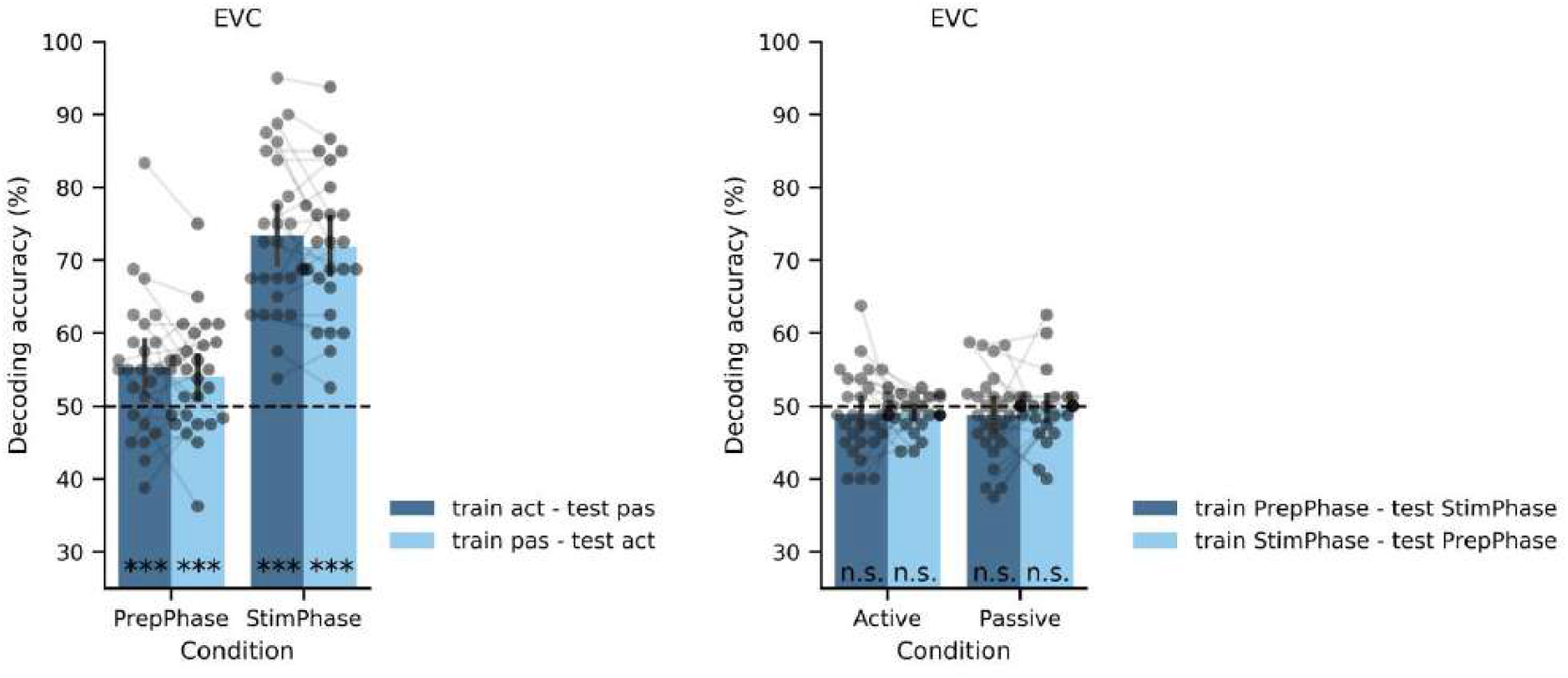
Results from multivariate analysis, cross-classification. Decoding accuracy is plotted for each single participant. The figure on the left shows results for training on active and testing on passive conditions (train act – test pas) and vice versa (train pas – test act). The figure on the right shows results for training on the preparatory phase and testing on the stimulus phase (train PrepPhase – test StimPhase) and vice versa (train StimPhase – test PrepPhase). Error bars represent the 95% confidence interval. Chance level is indicated by the dotted line at 50%. n.s. = not significant, *** = p < 0.001.

## Discussion

This study investigated how action-based predictions modulate activity in early visual cortex. Overall, our results do not support the cancellation, sharpening, and pre-activation accounts. Instead, our results mostly align with the opposing process theory. In line with our predictions for this theory, we show that both action-based and general sensory predictions induce similar stimulus-specific representations prior to stimulus presentation as evidenced by significant decoding and cross-decoding. During stimulus presentation we also found no differences in decoding accuracy between conditions, and cross-classification between conditions was possible. However, we observed enhanced – rather than suppressed – BOLD responses for self-generated stimuli, a finding inconsistent with all the aforementioned theories. Altogether, these results suggest that 1) action-based and general sensory predictions are based on similar mechanisms, and that 2) action execution can boost the amplitude, but not precision, of neural responses.

Our finding of enhanced rather than suppressed responses to self-generated stimuli challenges the forward model, also known as the cancellation account, which posits that efference-copy-based predictions suppress sensory consequences of actions. Although this theory has strong behavioral and neural support – showing reduced perceived intensity and neural responses for self-generated stimuli (Arikan et al., 2019; Bäß et al., 2008; Bays et al., 2005; S. Blakemore et al., 1999; S.-J. Blakemore et al., 1998; Cardoso-leite et al., 2010; Kilteni et al., 2020; Kilteni & Henrik Ehrsson, 2020; Lubinus et al., 2022; Martikainen et al., 2005; Straube et al., 2017; Voudouris & Fiehler, 2022; Weiss et al., 2011), we found no behavioral differences in orientation performance and an increased BOLD response for self-generated stimuli. Our findings align with other studies reporting neural enhancement (Csifcsák et al., n.d.; Krala et al., 2019; Reznik et al., 2014, 2015). It is still unclear why action sometimes suppresses and sometimes enhances neural activity. It is possible that the effect of action depends on task demands. In most studies showing suppression, participants had either no task at all, or had to judge the intensity of the stimuli. In the studies showing enhanced neural responses, participants had to closely monitor their actions, as they either had to play tones in the correct order or time (Reznik et al., 2014, 2015), or had to reproduce a specific distance traveled (Krala et al., 2019). Although our study did not require close action monitoring, participants performed a demanding discrimination task. Having control over when the stimuli would appear in the active condition may have increased attention and boosted neural responses in visual cortex. Variability across sensory modalities may also be relevant: suppression is more consistent in tactile and auditory domains, whereas visual studies report mixed effects. For example, while (Cardoso-leite et al., 2010) found reduced sensitivity for self-generated visual stimuli, (Schwarz et al., 2017) failed to replicate this result. Other behavioral studies show either enhancement or suppression depending on task timing (Desantis et al., 2014; Yon & Press, 2017). On the neural level, studies report reduced responses in visual areas (Benazet et al., 2016; Gentsch & Schütz-Bosbach, 2011; Lubinus et al., 2022; Ody, Straube, et al., 2023; Roussel et al., 2014; Straube et al., 2017), enhanced responses (Hughes & Waszak, 2011; Mifsud et al., 2016; Wen et al., 2018), or mixed effects depending on participants or ERP components (Buaron et al., 2020; Csifcsák et al., n.d.). Our study cannot determine the cause of the enhanced BOLD response in visual cortex for actively generated stimuli and the lack of behavioural differences. Future research should explore different tasks to test how task demands, sensory modalities, and stimulus predictability interact to drive suppression or enhancement of self-generated stimuli.

Our study also found no support for the pre-activation account, which proposes that learned action effects pre-activate sensory regions, resulting in early activation followed by perceptual suppression (Roussel et al., 2013). Several behavioural studies support this idea, showing early facilitation and later suppression of perception of self-generated stimuli (Desantis et al., 2014; Roussel et al., 2013; Yon & Press, 2017). Furthermore, it has been reported that active movements lead to an earlier BOLD response than passive movements (Kavroulakis et al., 2022). However, our univariate GLM could not find any significant differences between active and passive conditions in the preparatory phase. A deconvolution GLM revealed divergence just before stimulus onset, but with stronger sustained activity in the passive condition. This may either suggest a pre-activation in the passive condition, or a stronger initial dip in the active condition. If a stronger initial dip at the end of the prepatory phase is taken as evidence for more activity then there would be some indirect support to the pre-activation model. Multivoxel pattern analysis showed that the orientation of predicted upcoming gratings could be decoded from preparatory visual cortex activity. This finding aligns with previous research showing that predictions outside the context of action induce stimulus representations prior to their occurrence (Kok et al., 2017) or during omission of an expected stimulus (Kok et al., 2014). However, decoding performance was similar across conditions, and significant cross-classification suggests shared neural patterns. While EEG studies have reported stronger pre-activation for active conditions a few hundred milliseconds prior to action execution (Ody, Kircher, et al., 2023; Reznik et al., 2018), these short-lived effects may have been undetectable with fMRI. Overall, our results suggest that both action-based and general sensory predictions lead to similar pre-activation in early visual regions. This challenges the idea that action uniquely enhances pre-activation, highlighting the need for further investigation into the interplay between action, prediction, and neural dynamics.

The sharpening account was also not supported by our results. The concept of predictions leading to sharpening originates from the general perception literature. Kok et al. (2012) demonstrated that expected stimuli result in a reduced neural response yet improved decoding performance. This finding suggests that expectations sharpen the neural response, an effect that may appear as suppression when examined using univariate measures. (Yon et al., 2018) extended this analysis to self-generated stimuli and observed similar results: expected sensory action outcomes led to reduced activity but improved decoding compared to unexpected outcomes. These findings were taken as evidence that action does not suppress neural activity, but instead sharpens neural responses. However, we found no difference in decoding accuracy between active and passive conditions. Unlike Yon et al., who compared expected and unexpected action outcomes (all self-generated), we compared self-generated and externally generated stimuli, both of which were expected and temporally predictable. The absence of differences in decoding performance between conditions suggests that action-based and general predictions may operate similarly under predictable conditions. Since our study did not include unexpected events, we cannot confirm whether predictions lead to a sharpening of neural responses for expected compared to unexpected stimuli. However, our results indicate that action-based predictions do not sharpen neural responses more than general sensory predictions do.

Overall, our results largely align with the opposing process theory (Press et al., 2019). We observed pre-activation for both action-based and general predictions, and no difference in decoding accuracy during stimulus presentation – consistent with the theory’s prediction for expected stimuli. However, we did find enhanced BOLD responses for self-generated stimuli, which the theory does not fully explain in this context. One possibility is that nonspecific factors like increased attention or arousal associated with action boosted neural responses. Another explanation might be linked to motor-induced responses in the visual cortex as observed for example in mice (Fiser et al., 2016; Muzzu & Saleem 2021; Vasilevskaya et al., 2023; for an overview see Schneider, 2020). Our current design cannot rule out the influence of these additional processes. Future studies should manipulate the predictability of stimuli as well as attention to further disentangle these effects.

## Conclusion

Altogether, we have shown that action-based and general sensory predictions pre-activate neurons in visual cortex by inducing a representation of the predicted upcoming stimulus. There representations were similar for self-generated and externally generated stimuli, suggesting similar mechanisms. Furthermore, we have shown that action can boost the amplitude, but not precision, of neural responses to self-generated stimuli. These results mostly support the opposing process theory (Press et al., 2019).

## Acknowledgements

BvK received support from the Deutsche Forschungsgemeinschaft (DFG, grant KE 2016/2-1 and SFB/TRR 135, grant number 222641018, project A10). LFM has received funding for this project from the Biotechnology and Biological Science research Council (BBSRC) BB/V010956/1 (‘Layer-specific cortical feedback dynamics’). We thank Francis Crabbe and Veronika Penkova for their support in the fMRI data collection.

## Methods

### Participants

A total of 28 healthy, right-handed participants were invited to take part in the study, which was comprised of three sessions: one behavioural experiment, one fMRI experiment, and one fMRI session with retinotopic mapping and functional localisers needed to create individual regions of interest. All 28 participants completed the behavioural experiment (8 male, 20 female, mean age: 23.6 years, *SD* = 4.7). Furthermore, 26 participants additionally took part in the fMRI experiment, and 25 completed all three sessions. For the participant that took part in the fMRI experiment, but not the localiser session, we were able to generate regions of interest based on the main experiment. The final sample for the fMRI study thus included 26 participants (7 male, 19 female, mean age: 23.5 years, *SD* = 4.5). For three participants, one run each had to be excluded; one due to excessive head motion, one due to technical error causing button presses to not be recorded, and one due to the participant pressing the button at incorrect times in the active condition.

### Stimuli

The stimuli were sinusoidal gratings with 1.5 cycles per degree (cpd), presented at 50% contrast. They were presented in an annulus (outer diameter: 8° of visual angle, inner diameter: 0.5°) surrounding a black presentation dot of 0.3°, on a midgrey background. Auditory cues were pure tones of either 400 Hz or 1000 Hz, presented through MRI-compatible headphones. Stimuli were presented with PsychoPy 3.2.4 on a Windows 10 PC. The monitor had a refresh rate of 60 Hz.

### Experimental design

The behavioural and fMRI experiment used the same experimental design, with some small differences in timing. Each trial started with an auditory cue, presented for 300 ms, that predicted the overall orientation of the upcoming gratings. The mapping between the cue and the grating orientation was counterbalanced across participants, and trained in the behavioural experiment prior to the fMRI experiment. After the auditory cue, there was a preparation phase, during which only the fixation dot was present on the screen. This phase lasted three seconds in the behavioural experiment and eight seconds in the fMRI experiment. When the preparatory phase had passed, the fixation dot turned red for 300 ms. In active conditions, this served as a cue for the participant to press the button with their right index or middle finger, after which the gratings were presented immediately as a result of the button press. Participants were given a maximum of 3 s to press the button. In passive conditions, the fixation dot turning red signalled to participants that the gratings would be presented soon. The interval between fixation cue and presentation of gratings in the passive condition was the average time the participant took to press the button in active conditions in the training of the behavioural experiment. The two gratings were presented for 500 ms each, separated by an interstimulus interval of 500 ms. The orientation of the first grating was always +45° or −45° from vertical, depending on which auditory cue was presented at the start of the trial, and the second grating deviated slightly from this orientation by one of five angles (± 0.5° - 6.5°). After the offset of the second grating, participants were asked to judge whether the second grating was oriented clockwise or counterclockwise with respect to the first grating. A green fixation dot indicated the interval during which participants could respond (behavioural: until they answered, fMRI: max. 2 s). They indicated their answer using MRI-compatible buttons (clockwise = left index finger, counterclockwise = left middle finger). In the behavioural experiment, the next trial started after an intertrial interval of 2.5 seconds. In the fMRI experiment, the next trial started after a jittered intertrial interval, with a minimum mean duration of 9 s (range: 7-11) to which remaining time from the button press intervals were added.

### Procedure

Participants were first invited to the behavioural experiment. Here, they first trained the mapping between auditory cue, button, and grating orientation, and were then familiarised with the task. They then performed a training session with the main experimental paradigm, in which only the largest angle difference (6.5°) was used, and feedback on their performance was given. If performance was below 75%, this training session was repeated. After training, all participants reached at least 75% accuracy and were able to proceed with the main experiment. The fMRI experiment took part on another day. At the start of this session, participants performed a brief training session outside the scanner to refresh their memory. They then proceeded with the fMRI experiment inside the scanner. On a third day, retinotopic mapping and functional localiser data were collected.

### MRI acquisition

Functional MRI data were acquired using a 3T TIM Trio scanner (Siemens, Erlangen, Germany), using a 32-channel head coil. A multiband sequence was used to collect whole-brain functional data with the following settings: 45 slices interleaved; field of view [FoV] = 222 mm; repetition time [TR] = 1000 ms; echo time [TE] = 36 ms; flip angle = 60°; PAT mode = GRAPPA, multiband acceleration factor = 3, slice thickness = 3.0 mm, distance factor = 10%, and voxel size = 3 x 3 x 3 mm. A total of 995 volumes were collected per run in the main experiment, 800 volumes for the polar angle retinotopic mapping, and 196 volumes for the functional localiser. A T1-weighted anatomical scan (MPRAGE) was collected at the end of each scanning session.

### MRI analysis

fMRI data were analysed using BrainVoyager 22.0 for Linux and custom Matlab scripts in Matlab R2019b. Data preprocessing included slice time correction, motion correction, high-pass temporal filtering, and coregistration to each participant’s anatomical image. Both the anatomical and functional data were then normalised to MNI space. For the univariate analysis, functional data were additionally smoothed with 8mm. For the multivariate analysis, the unsmoothed data were used. A GLM was created with predictors modelling the two different time points of interest for each condition (active/passive and leftward/rightward gratings). The preparatory phase was modeled from the onset of the auditory cue to the end of the preparatory phase, and the stimulus phase from the onset of the first stimulus until the offset of the second stimulus. An additional predictor was included to model the time during which participants could answer in response to the orientation discrimination task, from the onset until the offset of the green fixation dot. For the univariate analysis, each predictor contained all trials of that condition, leading to a total of 9 predictors per run. For the multivariate analysis, each trial was modelled with a separate predictor, leading to a total of 81 predictors per run.

### VOI definition

Visual areas V1, V2, and V3 were defined using standard retinotopic mapping procedures based on the polar angle sequence. An additional region called „early visual cortex” (EVC) was created by combining V1/2/3. Within these regions, voxels responsive to our stimuli were selected using our functional localiser (t-contrast stimulus > fixation, p < 0.01 uncorrected).

### Univariate analysis

A VOI GLM was run on our regions of interest using the predictors described in *MRI analysis*. Enhancement for active trials in the preparatory phase was tested with the contrast (Active prep > Passive prep). We tested for suppression for active trials in the stimulus phase with the contrast (Passive stim > Active stim). An exploratory whole-brain univariate analysis was performed in addition using the same contrasts (see Supplementary Material).

### MVPA

MVPA was performed with custom Matlab scripts, implementing the LibSVM software (http://www.csie.ntu.edu.tw/wcjlin/libsv). A linear support vector machine was trained to discriminate leftward from rightward gratings using the beta images resulting from the GLM. Classifier performance was tested using a leave-one-run-out cross-validation approach, and permutation testing was performed to determine statistical significance. MVPA was performed separately for active and passive conditions, as well as preparatory and stimulus phases. Paired sample t-tests were performed to determine differences in decoding accuracy between active and passive conditions, separately for the preparatory and stimulus phases. In addition, cross-classification analyses were performed. First, the classifier was trained on active trials and tested on passive trials, and vice versa. Second, the classifier was training on the preparatory phase and tested on the stimulus phase, and vice versa.

### Eye tracking analysis

Eye movements were recorded with an MR-compatible EyeLink 1000 eyetracker system (SR Research, Osgoode, ON, Canada). We analysed the gaze coordinates during the preparatory and stimulus phase for active and passive conditions separately. The data were first smoothed with a smoothing window of 50 ms after which they were detrended and mean-corrected to remove linear drifts and correct for calibration inaccuracies. We then defined a fixation region-of-interest by a circle around the center of the screen with a radius of 1.5° and calculated the percentage of samples in which participants looked outside of this fixation area. We used paired-sample t-tests to determine whether the number of samples outside of the fixation area differed between active and passive conditions.

## Supplementary material

### Results

#### Univariate analyses - ROI

Univariate results for each individual visual region depicted in Fig. 1 resembled the findings found in our early visual cortex ROI reported in the main manuscript. There were no significant differences between active and passive conditions during the preparatory phase in any of the visual areas (V1: t(25) = −1.634, p = 0.115; V2: t(25) = −1.425, p = 0.167; V3: t(25) = −1.034, p = 0.311). During the stimulus phase, there was significantly more activity in the active compared to the passive condition in all visual areas (V1: t(25) = −4.445, p = 0.000157; V2: t(25) = −4.439, p = 0.000159; V3: t(25) = −4.355, p = 0.000198).

**Fig. 1.**
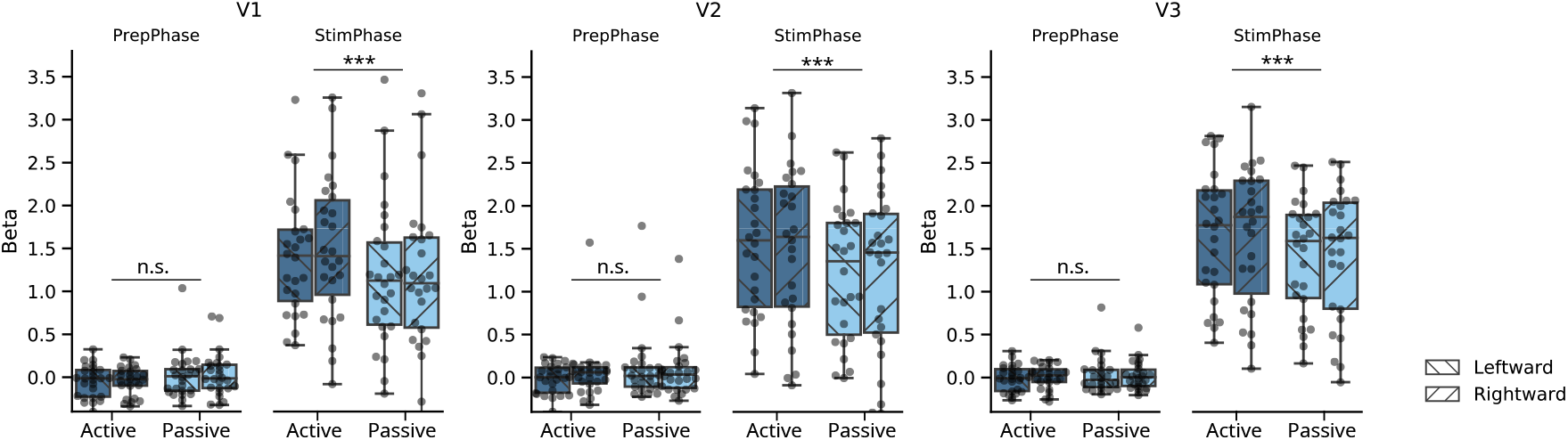
Univariate ROI analysis results. We performed paired t-tests to test for differences between active and passive conditions. Boxplots show the extracted beta values from V1, V2, and V3. n.s. = not significant, *** = p < 0.001.

#### Univariate analyses – Whole-brain

Whole-brain univariate analysis showed significantly more activity for active compared to passive conditions during the preparatory phase in supplementary motor area, left primary motor cortex, and right cerebellum (Fig. 2A). These areas are typically involved in motor preparation and therefore expected, as participants were preparing to press a button with their right hand in active conditions during this phase. Aligning with our ROI analysis, no significant differences were found in visual cortex. During the stimulus phase, we observed significantly more activity for active conditions in a large network with peaks in the cerebellum, supplementary motor area, primary motor and somatosensory cortex, middle and inferior frontal gyrus, insula, and precuneus (Fig. 2B). As participants pressed a button in active conditions during this phase, it is no surprise that many areas related to motor execution were observed. Supporting our ROI analysis, active conditions also enhanced activity in visual areas.

**Fig. 2.**
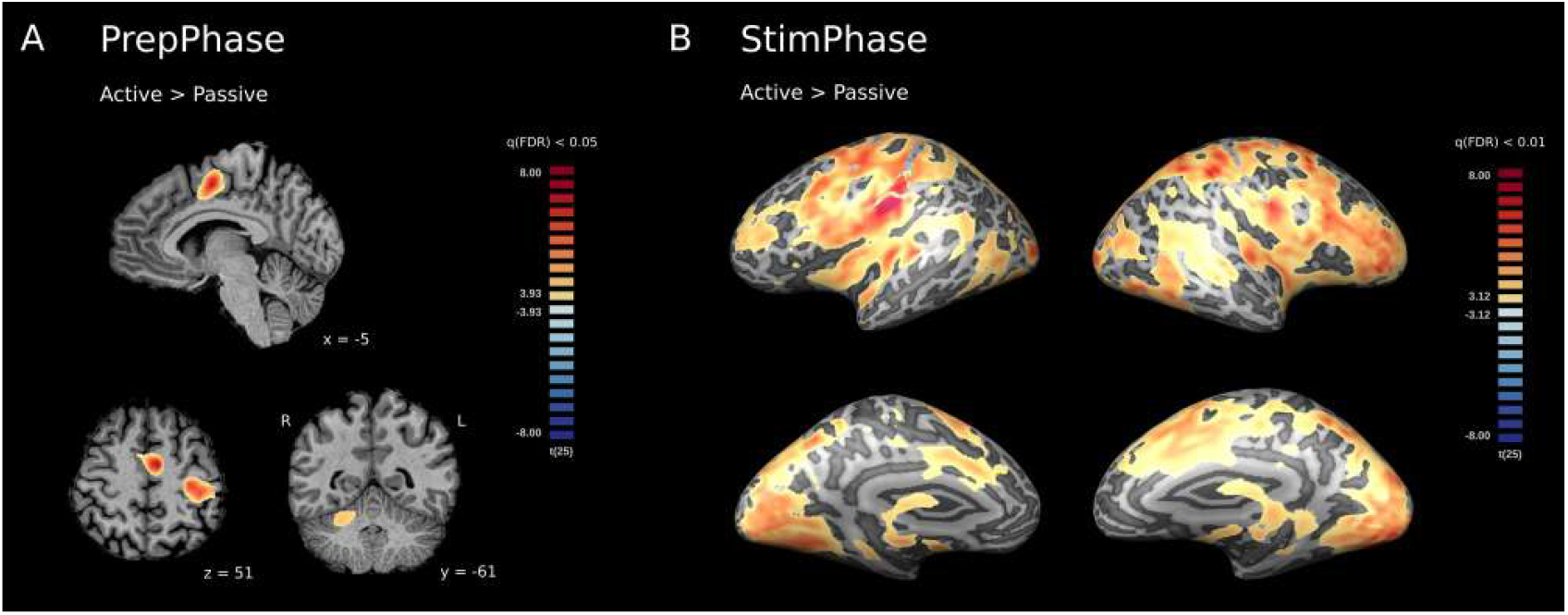
Univariate whole-brain analysis results. Active conditions were contrasted with passive conditions for the preparatory phase (PrepPhase, panel A) and for the stimulus phase (StimPhase, panel B). FDR: false discovery rate.

#### Multivariate analyses – Cross-classification

Cross-classification results for each individual visual region (Fig. 3) resembled the findings found in our early visual cortex ROI reported in the main manuscript. Training the classifier on active trials and testing on passive trials (and vice versa) showed significant above-chance decoding in both the preparatory and stimulus phase, indicating the classifier could generalise across conditions (Training on active and testing on passive trials, PrepPhase: V1 52.5%, p = 0.011, V2 54.2%, p = 0.002, V3 53.0%, p = 0.006; StimPhase: V1 67.5%, p < 0.001, V2 69.7%, p < 0.001, V3 67.9%, p < 0.001. Training on passive and testing on active trials, PrepPhase: V1 54.4%, p < 0.001, V2 54.2%, p< 0.001, V3 53.0%, p = 0.005; StimPhase: V1 66.6%, p < 0.001, V2 69.7%, p < 0.001, V3 64.8%, p < 0.001).

**Fig. 3.**
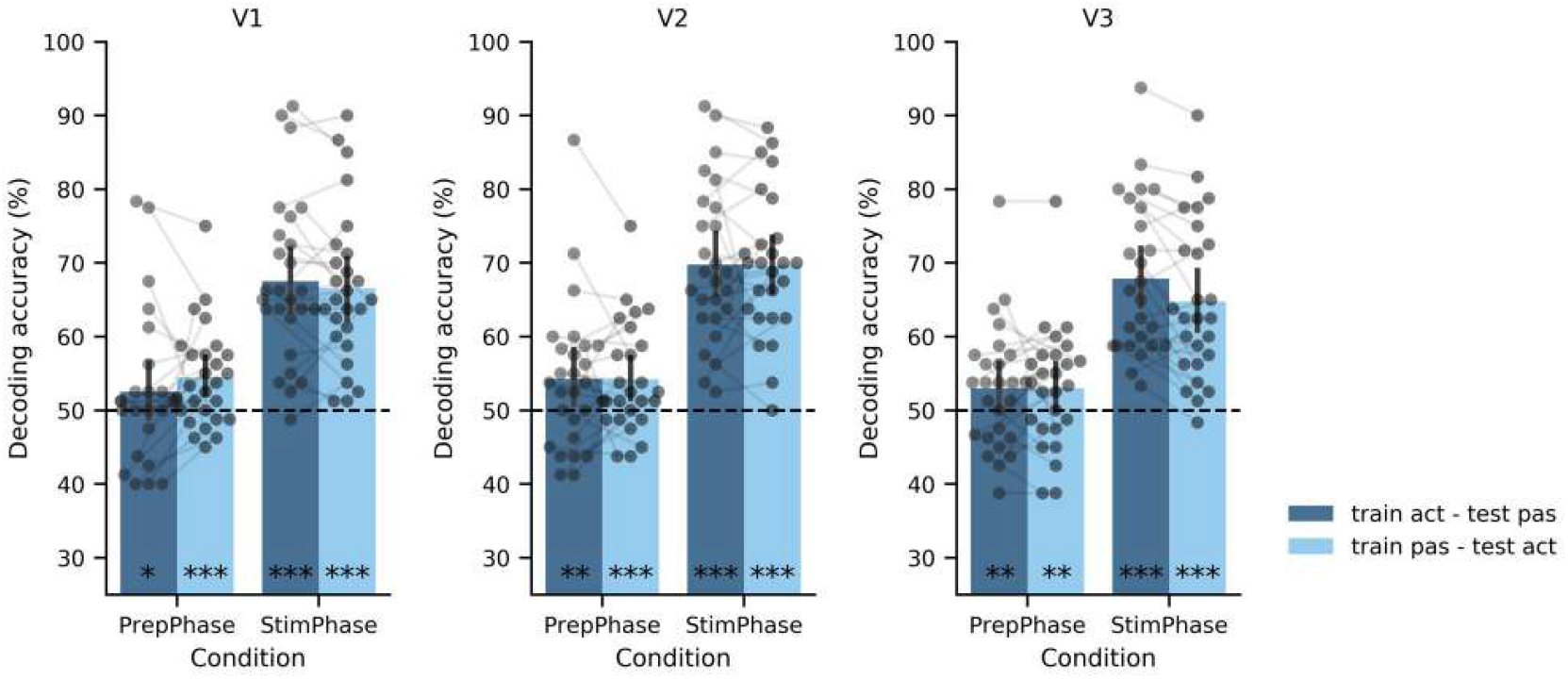
Results from multivariate analysis, cross-classification, for areas V1, V2, and V3. These are the results for training on active and testing on passive conditions (train act – test pas) and vice versa (train pas – test act). Decoding accuracy is plotted for each single participant. Error bars represent the 95% confidence interval. Chance level is indicated by the dotted line at 50%. * = p < 0.05, ** = p < 0.01, *** = p < 0.001.

In contrast, training on the preparatory phase and testing on the stimulus phase (and vice versa) showed chance-level decoding for both active and passive conditions, suggesting that the underlying patterns differed between these time points (Training on PrepPhase and testing on StimPhase, active conditions: V1 47.5%, p = 0.998, V2 48.7%, p = 0.902, V3 48.7%, p = 0.915; passive conditions: V1 48.6%, p = 0.910, V2 49.8%, p = 0.584, V3 48.4%, p = 0.938. Training on StimPhase and testing on PrepPhase, active conditions: V1 49.2%, p = 0.904, V2 49.6%, p = 0.778, V3 49.3%, p = 0.839; passive conditions: V1 51.1%, p = 0.03, V2 49.5%, p = 0.759, V3 49.3%, p = 0.871).

**Fig. 4.**
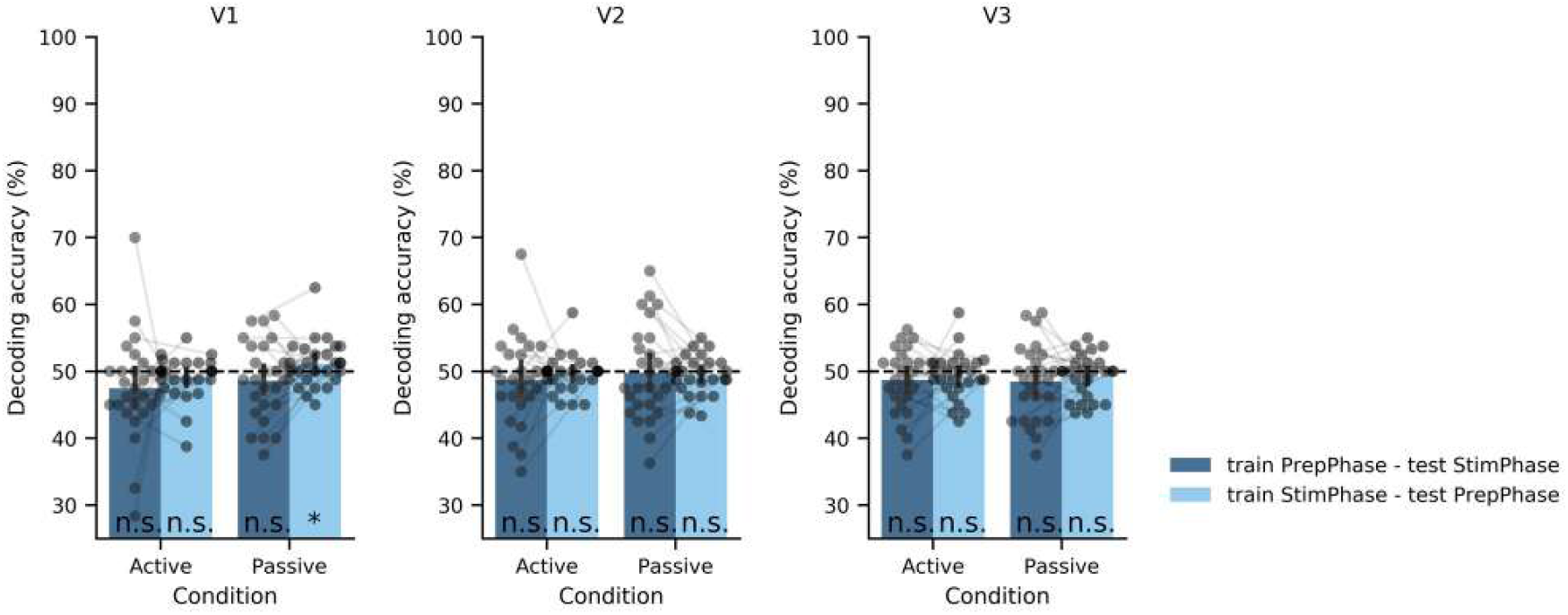
Results from multivariate analysis, cross-classification, for areas V1, V2, and V3. These are the results for training on the preparatory phase and testing on the stimulus phase (train PrepPhase – test StimPhase) and vice versa (train StimPhase – test PrepPhase). Decoding accuracy is plotted for each single participant. Error bars represent the 95% confidence interval. Chance level is indicated by the dotted line at 50%. n.s. = not significant, * = p < 0.05.

